# Measuring chromosome conformation by fluorescence microscopy

**DOI:** 10.1101/798512

**Authors:** Brian C. Ross, Fabio Anaclerio, Nicola Lorusso, Mario Ventura, Jim Costello

## Abstract

Measurement of *in-vivo* chromosome conformations (structures) in single cells is a major technological goal of structural biology. If one could identify many genetic loci in a microscope image despite the limited palette of fluorescent colors used to label them, then the conformation could be solved at some resolution by ‘connecting the dots’. Computational tools for making this reconstruction are expected to produce near-perfect reconstructions when the number of fluorescent colors is high enough, irrespective of the number of loci assayed. Here we report the first experimental test of the performance of these reconstruction algorithms and check their ability to reconstruct experimentally-measured conformations. We also demonstrate the experimental metrics needed to assess reconstruction quality. Our results indicate that current sequential FISH experiments may be close to the point where the reconstructions are nearly flawless at some distance scales.

---

The *in-vivo* conformation of chromosomes plays an major role in controlling gene expression [1] and is therefore of great importance to structural biology. Unfortunately there is not yet a direct assay of chromosome conformation in single cells. Until recently, most conformational inferences have been based on contact frequencies between pairs of chromosomal loci as measured by various chromosome-capture methods [2–7]. There is some doubt as to how well contact-based inferences work [8, 9], particularly when the contact frequencies have been averaged over many cells owing to high cell-to-cell variability [10, 11] (single-cell chromosome-capture is possible but technically difficult). Alternatively, fluorescence microscopy can directly measure the locations of a set of chromosomal loci, and ‘connecting the dots’ produces a conformation at the resolution of the locus spacing. The microscopy approach is becoming popular due to several advantages: it can directly measure the spatial locations of fluorophore-labeled chromosomal loci; like chromosome-capture it can assay many chromosomal loci in parallel; and unlike chromosome-capture it is inherently a single-cell method. However, its major weakness is scalability in number of loci: the number of genetic loci that can be identified in an image is usually taken to be equal to the number of different fluorophores that can be distinguished by color, typically ∼ 3 – 5. Recent developments in sequential FISH techniques [12] have pushed the number of uniquely identifiable loci beyond the number of colors, to around 50 in a typical experiment. However, a full measurement of chromosome conformation that resolves kilobase-scale structures will require thousands of loci to be discriminated, which is not yet possible using purely experimental tools.

In order to scale further to the thousand-locus scale, one can use computational tools. This final boost involves labeling many times more loci than can be discriminated by color or round of hybridization, and then computationally disambiguating look-alike spots in the image in order to reconstruct the underlying conformation. This disambiguation works by requiring loci that are nearby on the genome to also be close in three-dimensional space. The nearness requirement is obvious on small scales: for example, two loci spaced by 100 bp can only be physically separated 30 nm or less without the DNA breaking. But such a relationship has also been observed over kilobase-to-megabase scales [13–15], implying that computational reconstructions should also work at these larger scales. Simple reconstructions can be done ‘by eye’ [16], but two computational tools have been developed to automate this process and quantify the uncertainty in large reconstructions: `align3d` [17, 18] and `ChromoTrace` [19]. These tools have been tested using simulations but not yet using real-world experimental data.

A major advantage of reconstruction algorithms is that once the point is reached where adding more loci does not affect their density in the image, reconstruction performance should plateau. In other words, at this point reconstructions can be scaled up indefinitely, with little performance penalty. In this regime, reconstruction quality seems to be determined mainly by the number of what we term ‘pseudocolors’, which we define as groups of loci that can be distinguished purely experimentally. In a simple imaging experiment, the number of pseudocolors is simply the number of different fluorophores available: an experiment labeling some loci with red fluorophores and others with green fluorophores has 2 pseudocolors, since the red spots in an image can be discriminated from the green spots but not the other red spots, and vice versa. In a sequential FISH experiment the number of pseudocolors is the number of fluorophores times the number of rounds of hybridization: for example if 2 fluorophores are used over 5 rounds of hybridization, then labeled loci can be grouped into 10 categories (pseudocolors) when considering both color and round of hybridization. Simulations indicate that if labeling experiments use enough pseudocolors, reconstruction algorithms produce near-perfect results irrespective of the size of the reconstruction [18, 19].^1^

Here we describe the first tests of computational reconstruction algorithms using real-world labeling data, including both blind tests of the reconstructions (where the answer is genuinely unknown) and tests using published conformations where we have added ambiguity using a recoloring scheme (so that the reconstructions could be checked). Our immediate goal was to determine the performance on actual, as opposed to simulated or inferred, chromosomal conformations, and to validate as the accuracy of the reconstruction quality metrics that are produced by these analyses. In particular, the recolored data sets let us begin to address the question of whether current experiments have sufficient pseudocolors to solve the reconstruction problem; however recoloring produces fewer pseudocolors than can be attained experimentally, so we only directly test experiments that are less than state-of-the-art. As a secondary result, these recolored data sets allowed us to accurately model the distance function between pairs of loci, which itself gives clues to the biology of chromosome structure.

## Results

### 10-locus experiment

For our first reconstruction experiment, we fluorescently labeled 10 genomic loci over a 4 Mb stretch of chromosome 4 in fixed human cells (GM12878), using one of 3 spectrally distinct fluorophores to hybridize each locus (see Figure 1A). The cells were then imaged to localize the fluorescent labels in 3 dimensions, as illustrated in Figure 1B. There were several identical-looking spots in each color channel of the image, and the only way to identify these spots with specific loci was to compare the distances between them with the likelihoods of finding such distances given the known spacing of genomic labels, a relationship we call the interlocus distance function. In this experiment, we did not measure the distance function directly, but rather fit a Gaussian chain model [20] to the relationship between mean spatial distance between loci and their genomic separation found in Ref. [13], as illustrated in Figure 1C. Using the labeling pattern, the spot locations in each image, and the distance function, the reconstruction algorithm `align3d` produced likelihoods of assigning each spot in an image to each possible labeled locus of the same color, shown schematically as circles in Figure 2A. These ‘mapping probabilities’ can be used to infer likely conformations (Figure 2B) in which the level of confidence in each spot assignment is known.

**Figure 1.**
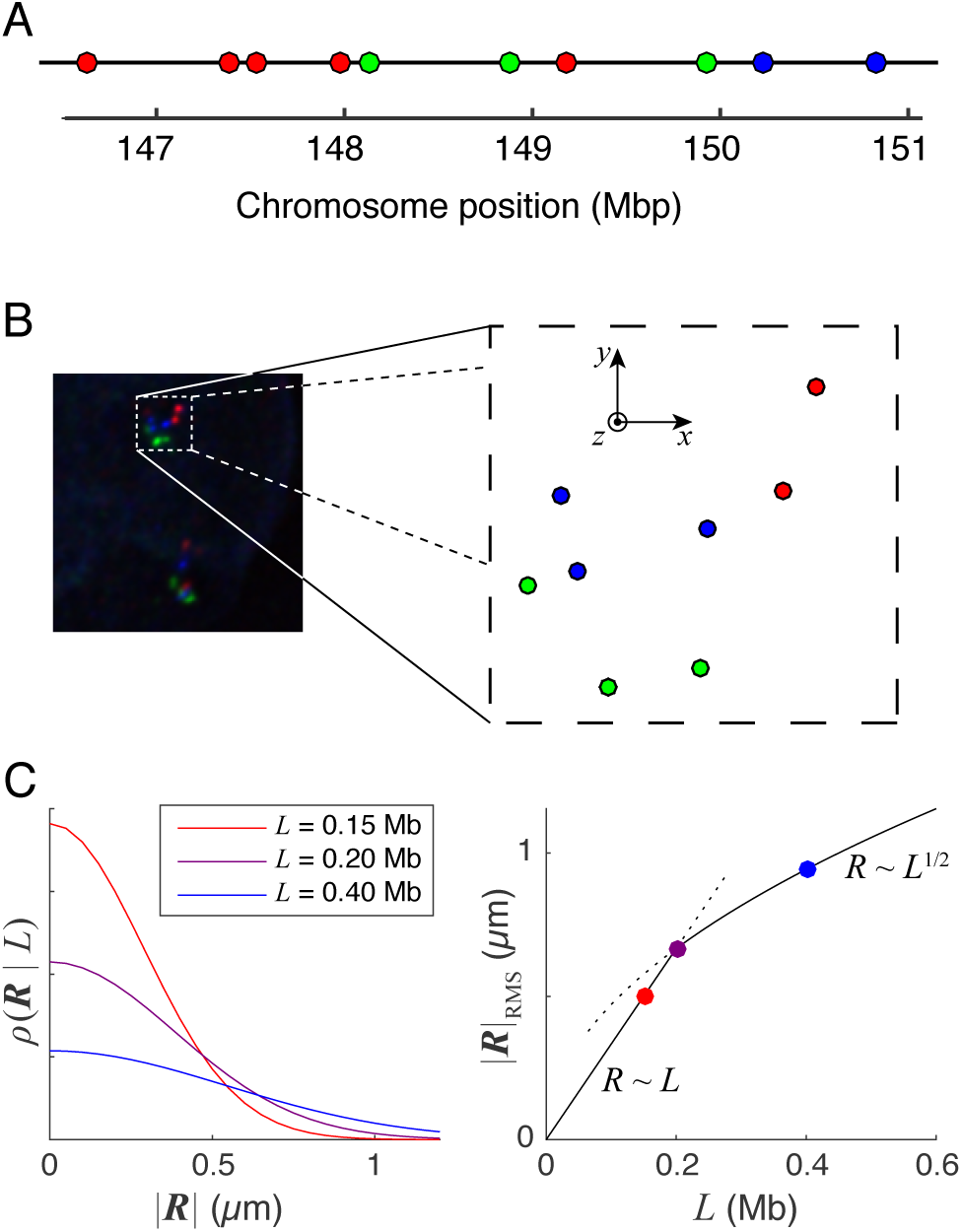
Inputs to the reconstruction algorithm. (A) Genetic locations and colors of fluorescent labels along human chromosome 4. (B) A summed z-stack of a typical cell image viewed from above, leading to a list of 3-dimensional spot positions and colors. (C) Left panel: distance function *ρ*(**R|***L*) modeled as a Gaussian for 3 values of *L*. Right panel: scaling of Gaussian width (RMS |**R**|) as a function of interlocus spacing *L*.

**Figure 2.**
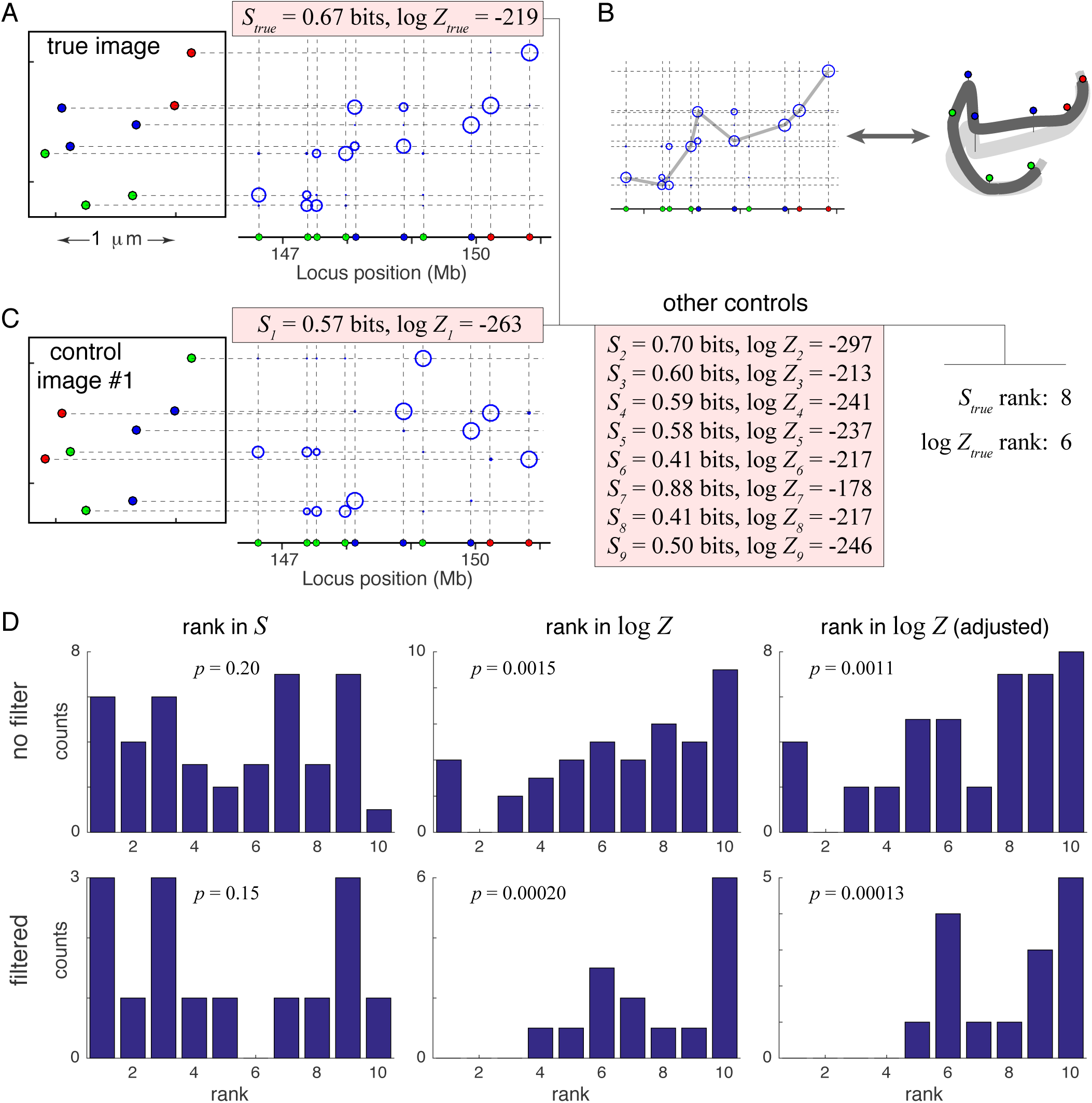
Reconstruction procedure (align3d). (A) `align3d` produces likelihoods of mapping the various labeled loci (*x* axis, at bottom) to the spots in the image (box at left), as shown schematically using circles. The uncertainty (entropy) *S* and the total statistical weight *Z* are outputs of this calculation. (B) By selecting a set of strong mappings one can infer a plausible conformation. (C) A control locus-to-spot mapping using a random permutation of spots colors. The rank-orderings of *S*_*true*_ and *Z*_*true*_ among the controls provide blind tests of the null hypothesis. *S* is expected to be lower, and *Z* higher, in the true mappings than the controls. (D) Distribution of ranks of the true *S* and *Z* over all experiments, without (top row) and with (bottom row) quality-filtering the experiments based on number of spots and chromosome thickness in *z*.

The true conformations were unknown, so the reconstruction quality could not be measured directly. However, we were able to test the null hypothesis of no significance by comparing each reconstruction with several ‘control’ reconstructions produced from identical inputs except that the imaged spot colors were randomly scrambled (see Figure 2C). Specifically, we compared two figures of merit between the true reconstruction and the controls: 1) the uncertainty in the spot assignments based on the mapping probabilities that we term ‘entropy’ (denoted by ‘*S*’), and 2) the total statistical weight over all possible conformations (denoted by *Z*). The expectation is that the entropy *S* is lower in the true reconstruction than the controls, and that *Z* (which roughly measures the ease with which the chromosome can be fit through the spots in the image) should be higher for the true mapping than the control mappings. We also looked at an adjusted *Z* that subtracts, in an approximate way, an unphysical contribution coming from the calculation (see Methods for details). The top row of Figure 2D shows the results of this test aggregated over all 42 chromosomes we measured. The bottom row shows the same tests applied to the 15 highest-quality chromosomes, selected as having 8-12 spots in the image and being more than 200 nm thick perpendicular to the slide (their width in the imaging plane was typically *∼* 1 *µ*m). In all metrics the true reconstructions outperformed the controls, as expected. *Z* was a more sensitive measure of significance than *S*, particularly when adjusted for artifacts of the calculation. Despite the much smaller sample size of high-quality chromosomes, the overall significance of their reconstructions against the null hypothesis was higher. Chromosomal thickness was an important determinant of quality (see Supplementary Figure S1), showing the importance of maintaining cell thickness with non-dehydrating fixation protocols in future experiments.

### Analysis of recolored published conformations

Our 10-spot experiments had two main limitations: 1) the distance function was not measured directly within the experiment, and 2) the reconstruction quality could not be checked directly since the true conformation was unknown (as would be the case in a practical use of our method, but ideally not in a method demonstration). To circumvent both these limitations, we reanalyzed two sequential FISH data sets that were published soon after our experiment, in which several tens of loci were uniquely identified and localized using variants of the protocol known as Oligopaints [12]. The first data set, which we refer to simply as the Oligopaints data [12], measured conformation at the megabase scale over whole chromosomes. The second data set, using the Oligopaints variant called ORCA [21], probed conformation at kilobase scales over small regions of chromosomes.

Oligopaints and ORCA are significant for us in three ways. First, these methods greatly increase the number of pseudocolors available to a reconstruction algorithm, potentially allowing the reconstruction of very large-scale conformations using hundreds or thousands of loci. Second, the interlocus distance functions could be measured precisely in Oligopaints data, since the locus positions were measured unambiguously. Third, by ‘recoloring’ the Oligopaints data to introduce ambiguity (see Figure 3), we could construct synthetic images requiring reconstruction, and then check the reconstruction quality using the known true conformations.

**Figure 3.**
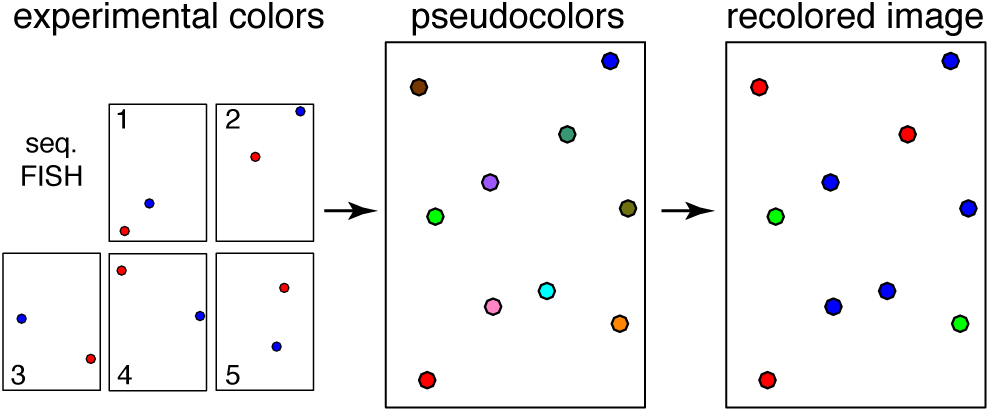
Recoloring procedure. Illustration of our recoloring scheme applied to the Oligopaints/ORCA sequential FISH data sets. Since these experiments uniquely localized each locus in a given image, the loci all have unique pseudocolors indicating their distinguishability. Our recoloring method randomly reassigns colors using a smaller palette (fewer pseudocolors than loci), simulates an experiment having more labeled loci than pseudocolors available. This process introduces ambiguity into the image, that requires a computational reconstruction to resolve. After applying our reconstruction algorithm to the recolored image, we can check the reconstruction quality using the original unambiguous locus assignments.

### Reconstructions at megabase scales (Oligopaints)

The first recolored data set we analyzed was that from the Oligopaints paper [12], which labeled TADs in several chromosomes of human cells. We were most interested in label spacing in the ∼ 0.5 Mb range, because at that scale adjacent loci should be near enough for reconstructions to work, but far enough that one should not need special superresolution techniques even when labeling many loci per pseudocolor. To obtain this rough label spacing, we restricted our analysis to the most densely-labeled parts of the Oligopaints data sets: one region in chromosomes 21 having 28 loci, and another region in chromosome 22 having 9 loci. Although the tiny chromosome 22 recolored conformations are probably not representative of future Oligopaints experiments, we think those reconstructions may be relevant for future live-cell experiments with limited pseudocolors.

Our first step was to use the Oligopaints data set to determine the interlocus distance function for these experiments and locus spacings. Our baseline assumption was that the distribution of locus separations would be described by the diffusive Gaussian chain model. However, we found somewhat more pairs of loci at large separations than predicted by the Gaussian chain model. This excess at large separations was well modeled by an exponential distribution in the interlocus separation *R*. As shown in Supplementary Figures S2 and S3, our distance function model was a sum of a Gaussian chain whose width scaled with interlocus distance according to an empirically-determined power law, and a exponential function that did not change with interlocus distance.

Next, we randomly recolored the Oligopaints data sets using 3, 5 and (for chromosome 21) 10 pseudocolors, and then passed the recolored images to `align3d` to obtain mapping probabilities for assigning spots to labeled loci. We also reconstructed 9 color-scrambled control images per ‘true’ recolored image, to compare the rank of the true image relative to the controls using various statistics of the reconstruction process. Note that the `align3d` calculation can only be done approximately for large calculations, so to simulate that we used a ‘baseline’ approximate calculation for all runs described here (see Methods section for details). Two statistics (entropy *S* and the adjusted statistical weight *Z*) proved sensitive in identifying the true reconstructions, particularly when 5 or more pseudocolors were used (see Supplementary Figures S4 and S5).

We measured the quality of the probabilistic `align3d` output using the metric of ‘unrecovered information’ *I*, which is the Shannon information required to specify the correct assignments given the locus-to-spot mapping probabilities. In the recolored experiments the true spot assignments were known, so *I* could be determined from the mapping probabilities, but in a real experiment one would approximate as *I* ≈ *S* since entropy *S* is simply a weighted average of information *I* over the mapping probabilities. A mapping that confidently assigns spots to their correct loci will have *S* ≈ *I* ≈ 0; if the spot assignments are very uncertain then *S* ≈ *I* ≈ log(*N/C*) for *N* loci per *C* pseudocolors; and if the spot assignment is confident but wrong then *S* is small but *I* → ∞. Figure 4 (left-hand panels) plots unrecovered information *I* versus entropy *S* over the various recolored Oligopaints reconstructions. These plots show that when the true mapping has significantly lower entropy than the controls, then *I* ≈ *S*, which is critical because it means that an experimenter can blindly gauge the quality of a reconstruction. We note that poor mappings send *I* ≫ *S*, probably because unrecovered information is very sensitive to errors: a single mistaken assignment of zero probability to a correct locus assignment sends *I* → ∞ but does not affect *S*. Next, we used each set of mapping probabilities to guess a likely conformation, using a heuristic formula that selects the most likely mapping probabilities and iteratively resolves conflicts where two loci map to the same spot. The right-hand panels of Figure 4 show the quality of the inferred conformations as measured by the rate of locus assignment errors, and show that this error rate can also be predicted by *S* although the relationship is weaker.

**Figure 4.**
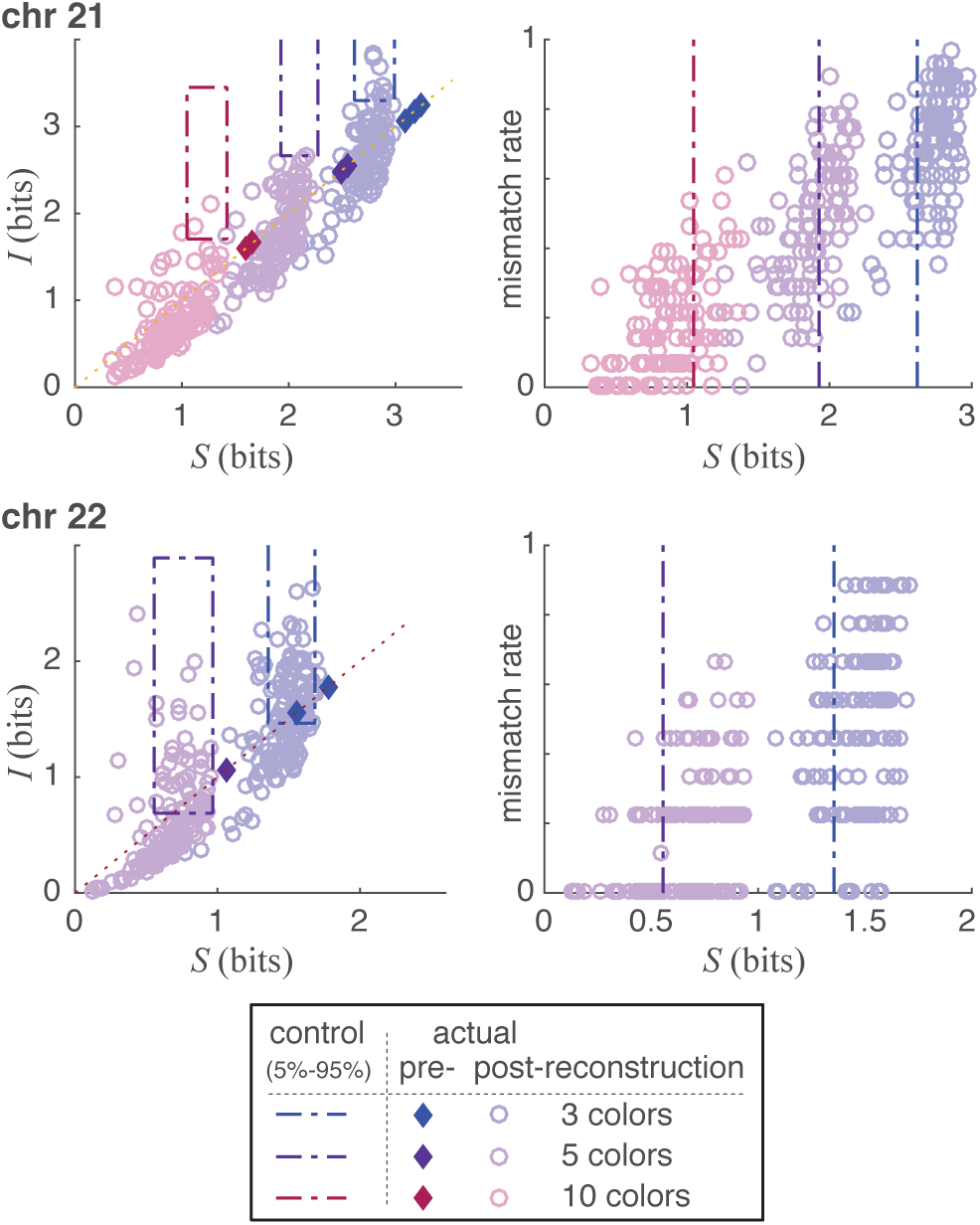
Reconstruction quality of Mb-scale (Oligopaints) conformations. (A) Actual information recovery *I* versus estimated information recovery *S*, for various recolorings of the Oligopaints conformations. Note that for these experiments we selected 28 closely-spaced loci from the chromosome 21 data set, and 9 closely-spaced loci from the chromosome 22 data set. The 5-95 percentile range of *I* and *S* from the color-scrambled (control) reconstructions fall within the respective dotted rectangles. (B) Rates of incorrectly-inferred locus identities as a function of estimated information recovery *S*. Dotted lines show the 5th percentile in *S* of their respective control: i.e. 95% of the controls had a higher value of *S*.

### Reconstructions at kilobase scales (ORCA)

Next, we tested our reconstruction performance using the fine-scale conformations found in the three ORCA data sets [21], whose labeled loci were spaced at 2, 3 and 10 kb respectively. Since the locus spacing was uniform we used the entire conformations for recoloring and reconstructions. First, we examined the distances between nearby loci to determine the interlocus distance function at small scales (see Supplementary Figures S6, S7 and S8). As before, the Gaussian chain model was adequate for the bulk of the distribution, but again a long tail indicated that nearby loci would take long excursions more often than a Gaussian chain would predict. As before, the residuals after subtracting the Gaussian chain distribution followed an exponential distribution. Thus we fit the 3 kb and 10 kb data sets using a Gaussian-plus-exponential distribution. At the smallest separations a second exponential was needed to provide a good fit.

We randomly recolored each of the three ORCA data sets using 3, 5, 10 and 20 pseudocolors, produced locus-to-spot mapping probabilities using `align3d` and inferred explicit conformations using those mapping probabilities, as before. The quality of the mapping probabilities and the locus assignments of the explicit conformations are shown in Figure 5, and their rankings relative to controls are shown in Supplementary Figures S9, S10 and S11. As before, when the true reconstruction outperforms controls, the entropy *S* accurately measures both unrecovered information *I* and locus-assignment errors, which are closely-related error measures (see Supplementary Figure S12). These reconstructions performed noticeably worse than the Oligopaints reconstructions, as measured by the greater values of unrecovered information *I* and error rates in the conformations for the same number of pseudocolors. There are several likely reasons for this. 1) The recolored ORCA data had more labeled loci than the recolored Oligopaints data. 2) The regular spacing of the ORCA labels made reconstruction more difficult. (In the Oligopaints data sets, irregularity in label spacing and color ordering are the two characteristics that help identify genetic loci). 3) The occasional long-distance excursions taken by loci (corresponding to the exponential tail in the distance function) are more significant at small scales. The farther these excursions go relative to a typical displacement between two adjacent loci, the more difficult reconstructions become, since the volume where neighboring loci may be found grows to encompass many more competing loci. In support of this last explanation, we note that the 3 kb and 10 kb data sets gave better results than the 2 kb data set.

**Figure 5.**
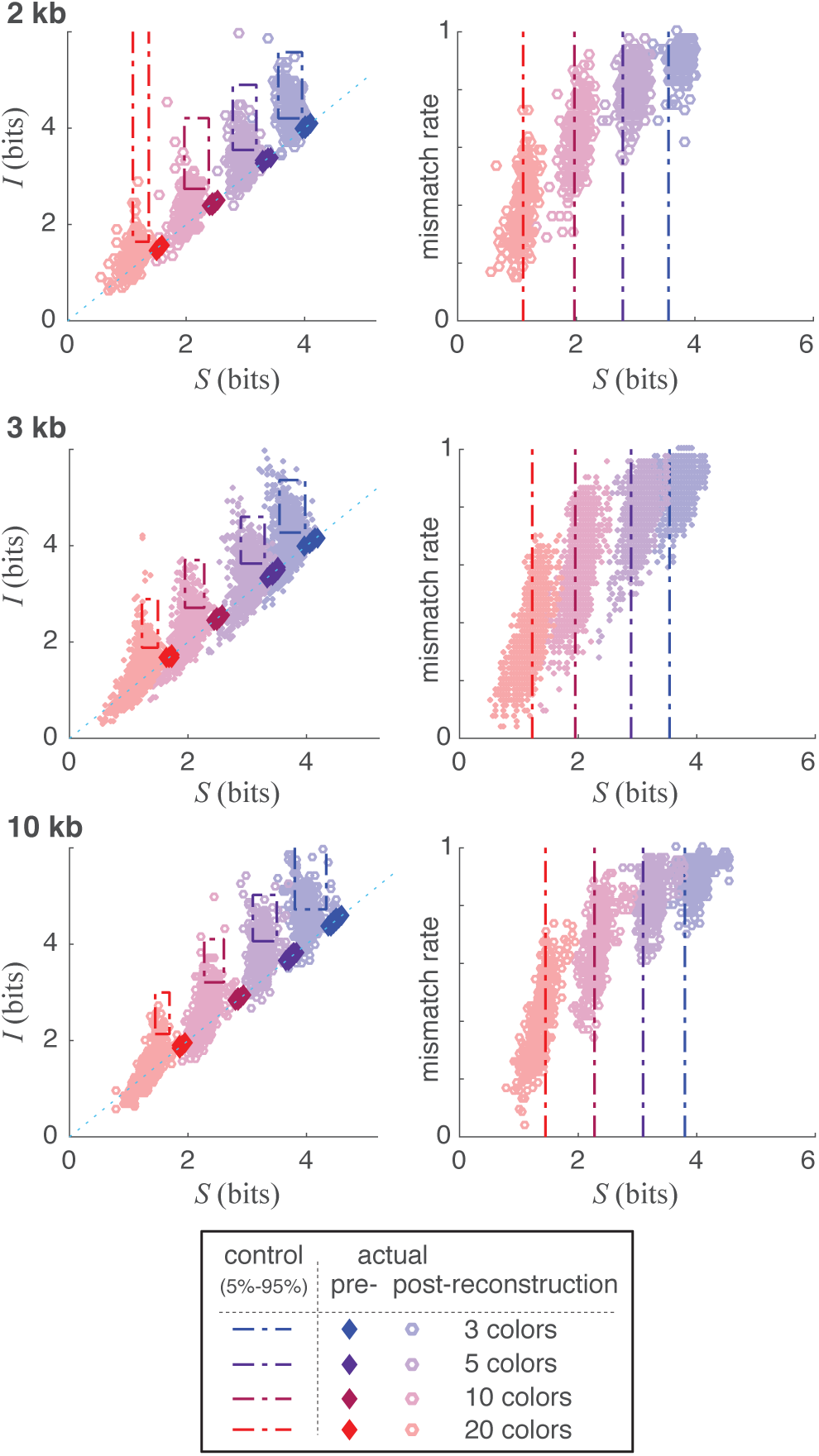
Reconstruction on kb-scale (ORCA) conformations. (A) Actual information recovery *I* versus estimated information recovery *S*, for various recolorings of the ORCA conformations. Dotted rectangles show the 5-95 percentile range of *I* and *S* from the respective control mappings. (B) Rates of incorrectly-inferred locus identities as a function of estimated information recovery *S*. Dotted lines show the value of *S* lying below 95% of the values of *S* from color-scrambled control reconstructions.

## Discussion

Conformation capture experiments such as Hi-C provide much useful information about genome architecture, but they are not an ideal tool for measuring explicit chromosomal conformations, for several reasons. First, most Hi-C data comes from large cell populations whose conformations are likely very dissimilar: a ‘consensus conformation’ deduced from these experiments is mostly meaningless [8], especially given that the structures visible in population-averaged data only exist in a small fraction of cells [10, 11]. Single-cell chromosome-capture experiments are possible but they have lower resolution [10]. Second, these experiments require deep sequencing, which is expensive. Third, even single-cell contact-based structural inference has the overfitting problem where many continuous degrees of freedom are adjusted to fit noisy contact data: the resulting conformations may plausibly fit the data, but the uncertainty in these guesses is completely unknown. Finally, some important information about chromosomal positioning is inevitably absent from contact data, the most obvious examples being the location, orientation and ‘parity’ of the chromosomes.

By contrast, image-based chromosomal reconstructions avoid many of the problems of Hi-C reconstructions, as our results demonstrate. Fluorescence imaging uses cheap reagents, is widely available and is inherently a single-cell assay. The reconstruction method we demonstrate factors in the various types of experimental error. It produces definite conformations where the data is unambiguous, and gives the spectrum of possible conformations with accurate likelihoods where the reconstructions have uncertainty (even when the calculation is only approximate – see Supplementary Figure S14). The inference itself requires finding a finite set of discrete mapping variables, rather than a continuum of real-valued parameters as in the case of Hi-C reconstructions: thus it is far less underdetermined than Hi-C reconstructions, and its output is able to consider all possible solutions and properly weight their likelihoods. We believe our permutation tests are practically immune to known and unknown systematic effects when the label colors are chosen randomly. In turn, when a reconstruction is significant vis-á-vis the permutation tests, the specific error rate in the reconstruction can be accurately determined from the entropy *S*. For these reasons we believe that conformational reconstructions from microscopy have much better-understood error bounds than those from Hi-C data.

Refs. [18] and [19] argue that once experiments attain sufficient pseudocolors, the conformational inference problem should be essentially solved for reconstructions of any size, and that there is little purpose in improving experimental techniques further. The question is whether we have already reached the required number of pseudocolors with state-of-the-art sequential FISH protocols. We cannot yet give a complete answer from the recolored sequential FISH data, because while those methods are capable of ∼ 50 pseudocolors, the data sets localized only ∼ 50 loci so our recolored data sets necessarily used far fewer than 50 pseudocolors. A full experiment labeling hundreds or thousands of loci using the full 50 pseuodcolor palette could perform worse or better than our 20-pseudocolor results: the density of competing spots will increase (if for no other reason than the fraction of spots on the boundary will drop), but on the other hand the number of pseudocolors will be much higher. However, our 3- and 5-pseudocolor results should directly correspond to the reconstruction performance in live-cell experiments where only 3 − 5 pseudocolors are available. One result we are already confident in even for static experiments is that kilobase-scale reconstructions will require more pseuodcolors than megabase-scale reconstructions having the same number of loci, since the long tail in the distance function increases the search radius relatively much more when looking at small scales.

The interlocus distance functions themselves reflect interesting biology. Our models are consistent with chromosomal DNA taking a simple random walk polymers occasionally punctuated by long-distance excursions, having an exponentially-decaying likelihood with distance that reflects some unknown biology. One might expect that long stretches of DNA should undergo more excursions than short ones, but surprisingly the interlocus distance functions we fit show that the rate of excursions is *constant* no matter the length of intervening DNA. Our explanation is that the DNA is initially bundled as a random-walk polymer, and subsequent processes in the cell cause certain genomic regions to become stretched away from their initial position without affecting DNA on either side of a given affected region. Thus when measuring the positions of two loci, an excursion is only registered if one of the two loci happens to be in the part of DNA being stretched: interior stretches ‘leave and come back’ between the loci and are not registered. The presence of two exponential functions in the small-scale 2 kb ORCA data set is consistent with the fact that one or both endpoint loci can be undergoing an excursion, and the coarser data sets most likely only register excursions taken by both endpoints.

Several experimental lessons come out of our early results. First, having fully three-dimensional conformations is not only important for preserving genomic integrity, but also essential for obtaining accurate locus-to-spot inferences. In the flattest cells of our original experiment, the collapse of the out-of-plane dimension handicapped our analysis in the same way that simply ignoring the *z*-coordinate information in our spot data would. Second, it is important for the experimental color channels to be aligned, as this significantly affects the spacing between spots of different colors (and hence pseudocolors). Third, it is important to measure the interlocus distance function within each experiment, ideally by hybridizing set of probes labeling 1 locus per pseudocolor to a parallel preparation of cells. A good example showing why this is important is the difference in distance functions between the ORCA 2 kb and 10 kb data sets versus that of the ORCA 3 kb data set. Fourth, we find slight evidence that certain random labeling patterns are slightly better than others (see Supplementary Figure S13) though the effect is slight, but we did not look into nonrandom labeling patterns.

The computational protocols we developed here will be useful in the reconstruction of much longer and higher-resolution conformations. Computation time for all reconstructions in this paper including controls was only *∼* 3 processor-days, so if experiments can scale to assaying thousands of loci per cell, then the reconstructions easily can too. If the observed trend in quality versus number of pseudocolors extends to the regime of *∼* 50 pseudocolors, then large-scale conformational reconstructions are already possible using experimental techniques which are becoming routine.

## Methods

Interphase nuclei were obtained from lymphoblast cell lines (GM12878). FISH experiments were performed using human fosmids containing non-duplicated region directly labeled by nick-translation with Cy3-dUTP (PerkinElmer), Cy5-dUTP (PerkinElmer), and fluorescein-dUTP (Enzo) as described by Ref. [22], with minor modifications. Briefly, 300 ng of labeled probe were used for the FISH experiments; hybridization was performed at 37 ° C in 2x SSC, 50% (v/v) formamide, 10% (w/v) dextran sulphate, and 3 mg sonicated salmon sperm DNA, in a volume of 10 ml. High stringency post-hybridization washing was performed at 60 ° C in 0.1x SSC three times.

Nuclei were simultaneously DAPI-stained. Digital images for 2D-FISH were obtained using a Leica DMRXA2 epifluorescence microscope equipped with a cooled CCD camera (Princeton Instruments). DAPI, Cy3, Cy5, and fluorescein fluorescence signals, detected with specific filters, were recorded separately as grayscale images. Coloring and merging of images were performed using ImageJ. In total, 21 interphase cells were scored.

Spots were located and their x/y centers were recorded manually from the deconvolved image stack. (The positional error introduced by deconvolution was found to be about 30 nm.) The z position of each spot was found to sub-stack-spacing resolution by fitting the intensity profile of each spot’s x/y pixel over the image stack to a Gaussian. Each spot on the color-calibration slide was localized in all color channels, and the difference of each channel from the bead was used to build a linearly-interpolated chromatic error model. This model was used to correct the localizations of the single-color fluorophores on the 3-spot and 10-spot slides. Lastly, spots were grouped into 2 chromosomes on each of the 10-spot slides.

Locus-to-spot mapping probabilities for each chromosome were produced using the `align3d` tool [17], which is available for download at https://github.com/heltilda/align3d. For the 10-spot experiment we computed the mapping probabilities exactly (to within the accuracy of the distance function we provided), by enumerating each possible conformation consistent with the data and weighing it according to the distance function. For the recolored experiments, we wanted to simulate the performance of a much larger reconstruction where such enumeration is impossible, so we used the lowest-order approximation (denoted *Z* in Ref. [17] or *Z*0 in Ref. [18]) that optimized spot penalties to approximate the true mapping probabilities.

In the original `align3d` paper the total statistical weight *Z* contains an unphysical contribution from spot penalties and skipped-spot penalties, which are part of the calculation and unrelated to the ease of fitting physical contours through the spots. For purposes of computing rankings of true reconstructions versus controls, we considered an ‘adjusted *Z*’ which removes those factors in an approximate way by calculating the following quantity:

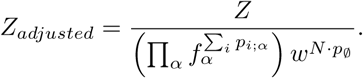

## Acknowledgments

We gratefully acknowledge support and helpful discussions from Andrew Laszlo, Joshua Vaughan, Leila Garcia, Min Fang, Julia Sidorova, Bob Kao, Jenny Mae Samson, Jens Gundlach, and Huy Nguyen. This work was supported by the Boettcher Foundation (J.C.), NIH grant 2T15LM009451 (B.R.), and a Cancer League of Colorado grant (B.R.).

## Supplementary Figures

**Supplementary Figure S1.**
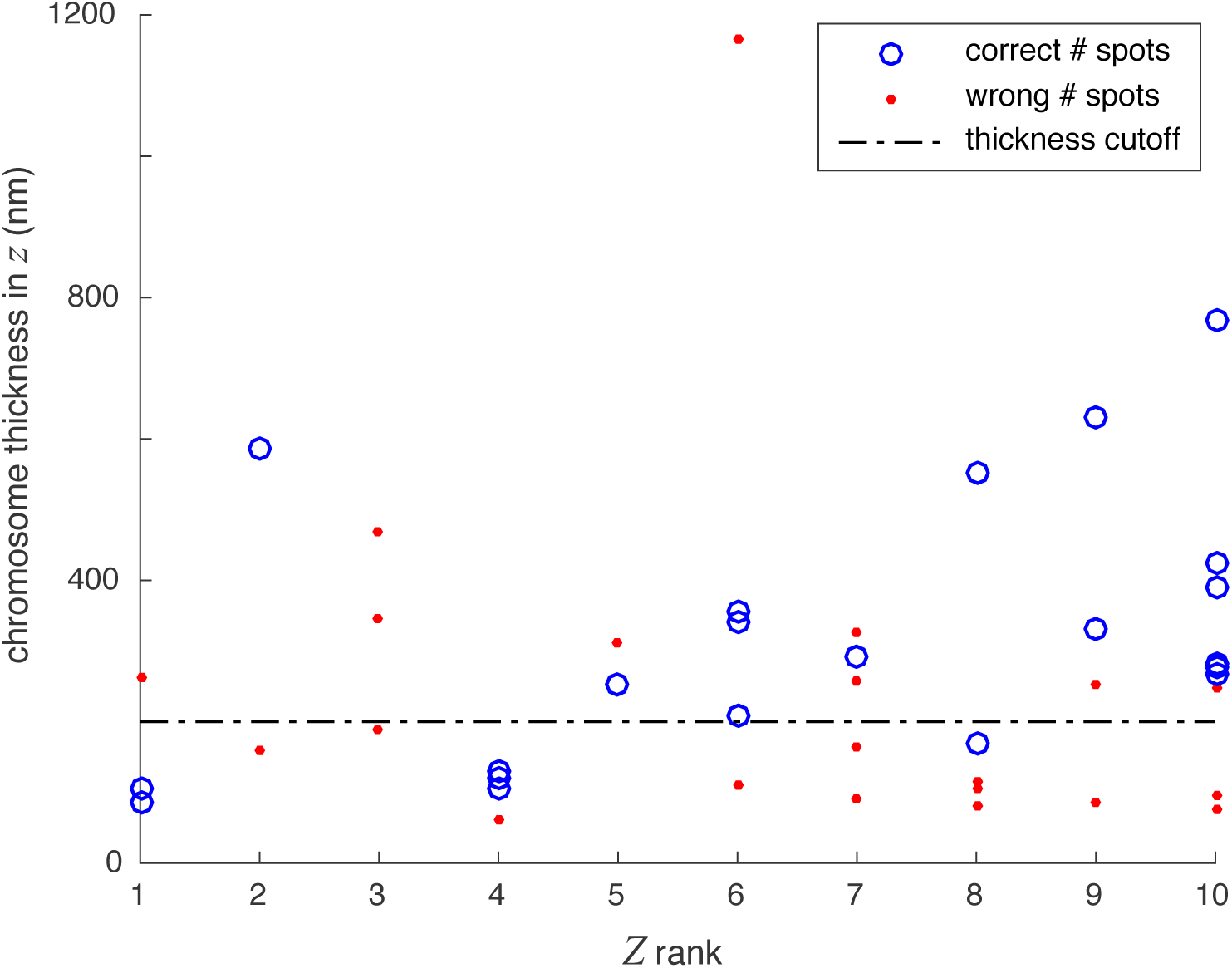
Chromosome thickness vs. rank in statistical weight *Z*. Thickness of chromosomes in *z*, plotted versus rank in the statistical weight *Z* of the true mapping versus controls. Thickness was measured from 2nd-lowest to 2nd-highest spot, to exclude outliers. Chromosomes with near-correct numbers of imaged spots (8-12 out of 10 expected) are plotted in blue; chromosomes with fewer than 8 or more than 12 imaged spots are plotted in red. Filtering the chromosomes on a) having 8-12 spots, and b) having reasonable thickness in *z* (greater than one 200 nm imaging slice) causes the average *Z* rank of the true mappings to increase relative to the controls, in line with the expectation that good mappings should have higher statistical weight than controls. The improved test against the null hypothesis indicates that preserving cell thickness is important for obtaining good reconstructions. Note that even the good chromosomes here are very flat in *z* compared with *x* and *y*, indicating that the ranks would probably have been higher yet if volume-preserving fixation were used.

**Supplementary Figure S2.**
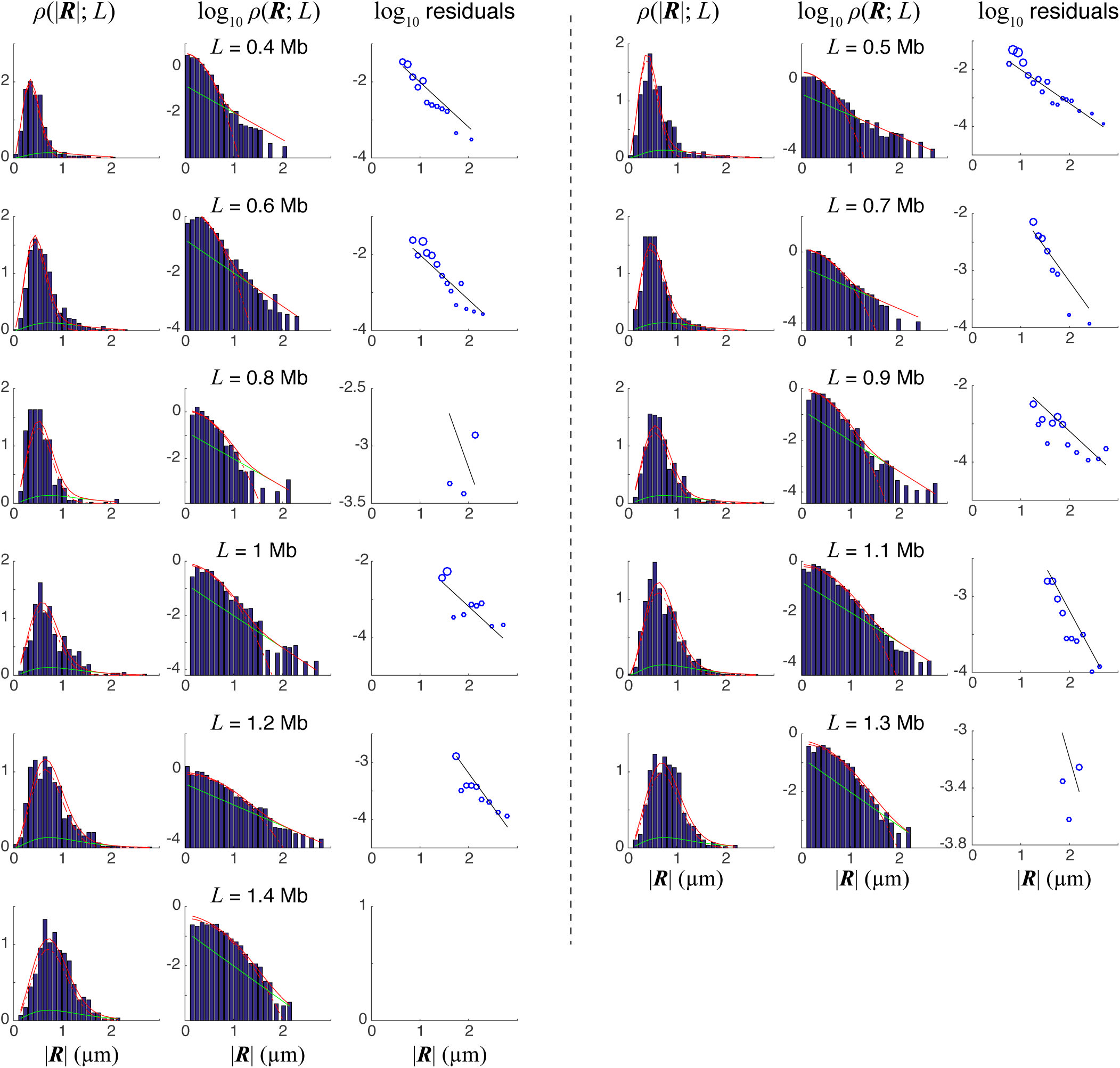
Distance function for Oligopaints chromosome 21 reconstructions. The observed probability distribution *ρ* of spatial distances between pairs of loci (blue histograms) overlaid with the model fit (solid red line), binned for different values of the interlocus distance *L*. We modeled *ρ* as a sum of a Gaussian chain distribution (*p*_*Gauss*_ = 0.81369, |**R**|_*RMS*_ = 0.0001303 *µ*m *L*^0.60817^ for *L* in bp; dotted red line) and an exponential decay in |**R**| (*p*_*exp*_ = 0.18631, decay constant 2.7209 bp^*−*1^; green line). First plot for each value of *L* shows the likelihood of observing a given separation distance; second plot shows the log distribution for observing a given separation vector **R**; third plot shows the residuals of log^1^0 *ρ*(**R**; *L*) when the Gaussian chain is subtracted from the observed distribution, overlaid with the model exponential capturing these residuals. There are no empty bins in the histograms, as they have been absorbed neighboring bins: therefore an ideal fit would touch the top of every bin.

**Supplementary Figure S3.**
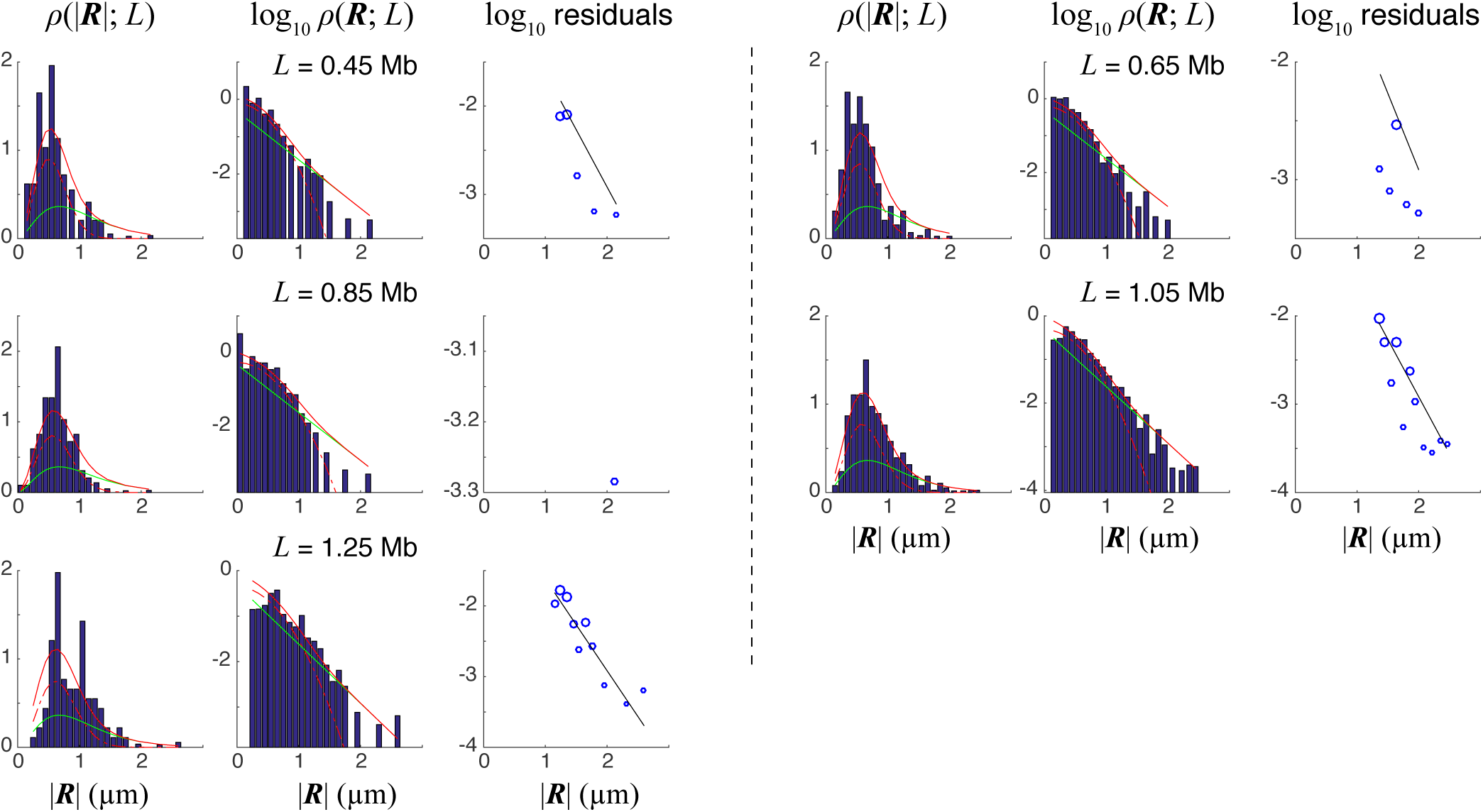
Distance function for Oligopaints chromosome 22 reconstructions. Observed probability distribution *ρ* of spatial distances between pairs of loci (blue histograms) overlaid with the model fit (solid red line), at different values of the interlocus distance *L*. See Figure S2 caption for details. We modeled *ρ* as a sum of a Gaussian chain distribution (*p*_*Gauss*_ = 0.54969, |**R**|_*RMS*_ = 0.042279 *µ*m *· L*^0.18933^ for *L* in bp; dotted red line) and an exponential decay in |**R**| (*p*_*exp*_ = 0.45031, decay constant 2.9893 bp^*−*1^; green line).

**Supplementary Figure S4.**
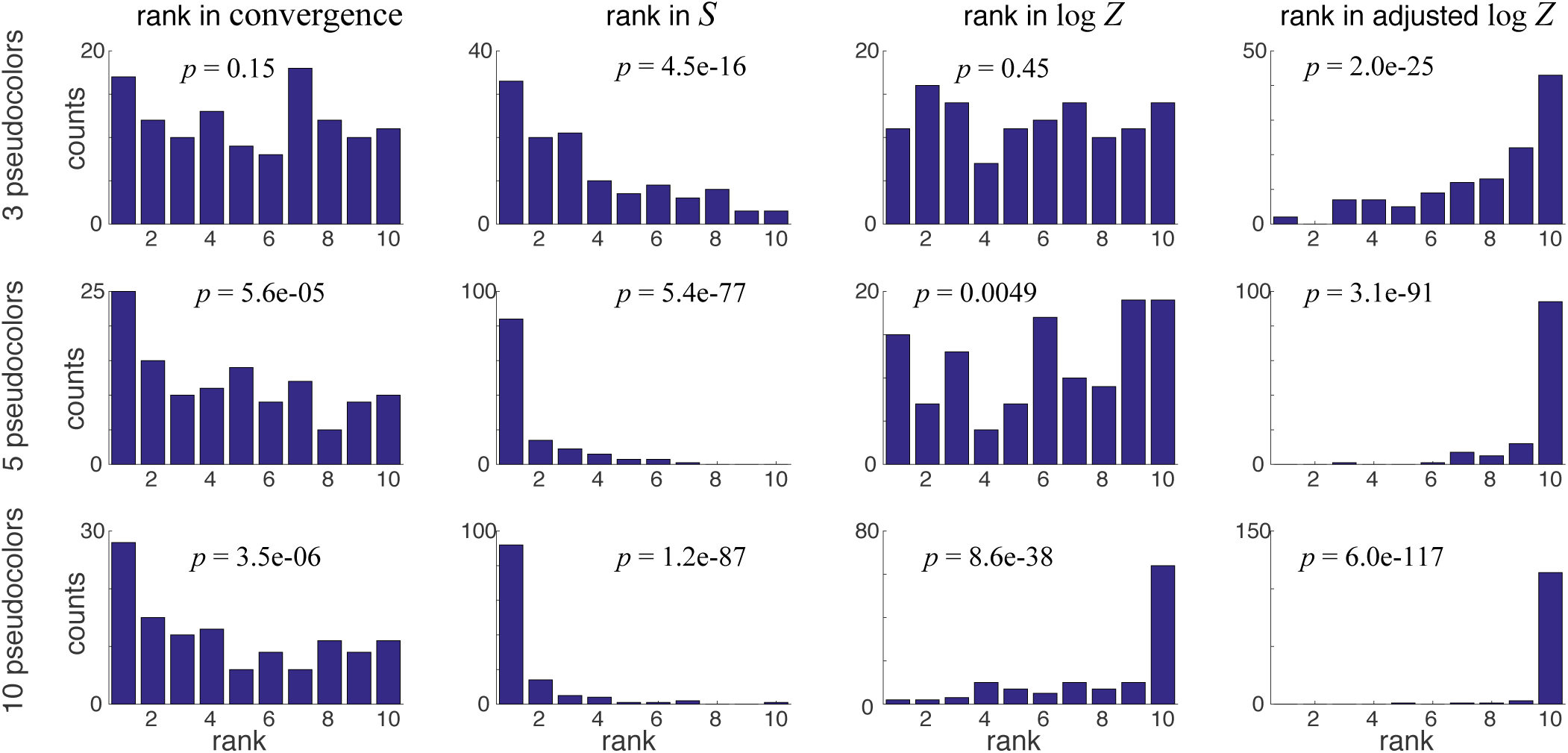
Null hypothesis tests of Oligopaints chromosome 21 reconstructions. Ranks of several statistics from the true mappings relative to those from control mappings, for three different recolorings. The statistics are: 1) the number of iterations required to converge the calculation; 2) the mapping entropy *S*, 3) the total weight *Z* (or equivalently its logarithm); 4) the total weight *Z* adjusted to approximately subtract the contribution coming from the unphysical free parameters. The expectation is that the true mapping will have: fewer iterations (lower rank in the convergence column), lower *S*, and higher log *Z* (with and without adjustment). *p*-values are the likelihood of randomly selecting aggregate ranks as low (first 2 columns) or as high (last 2 columns) as the aggregate (summed) ranks observed.

**Supplementary Figure S5.**
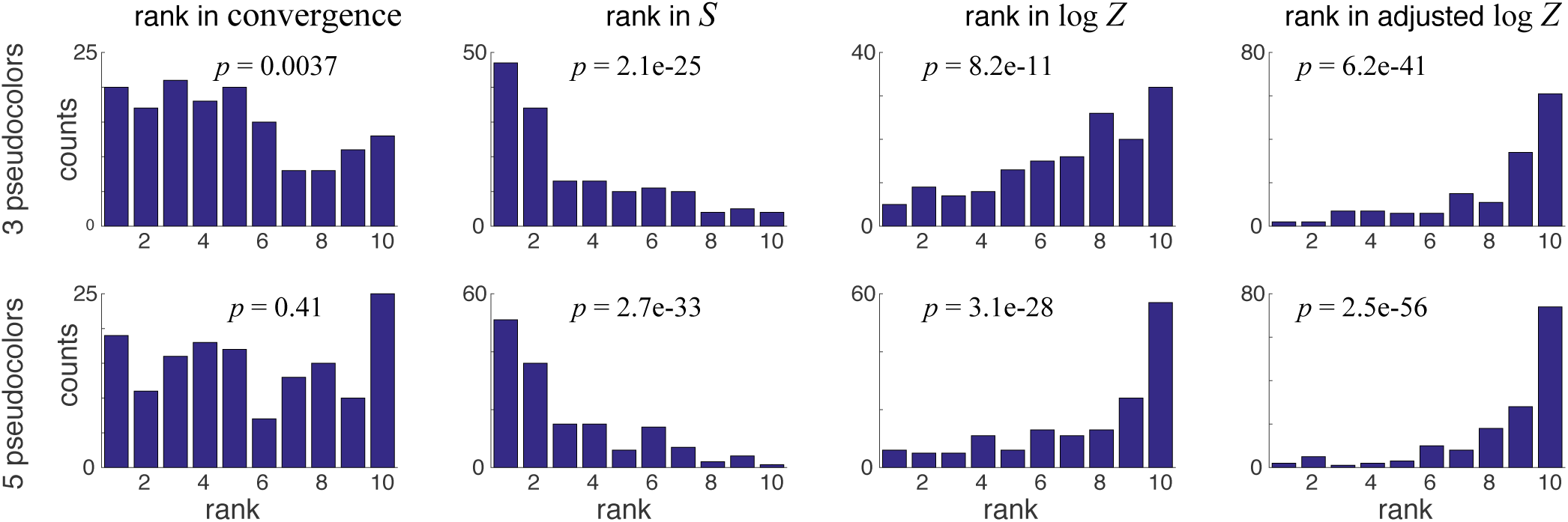
Null hypothesis tests of Oligopaints chromosome 22 reconstructions. Ranks of mapping statistics from the true chromosome 22 mappings relative to the controls. See Figure S4 caption for details.

**Supplementary Figure S6.**
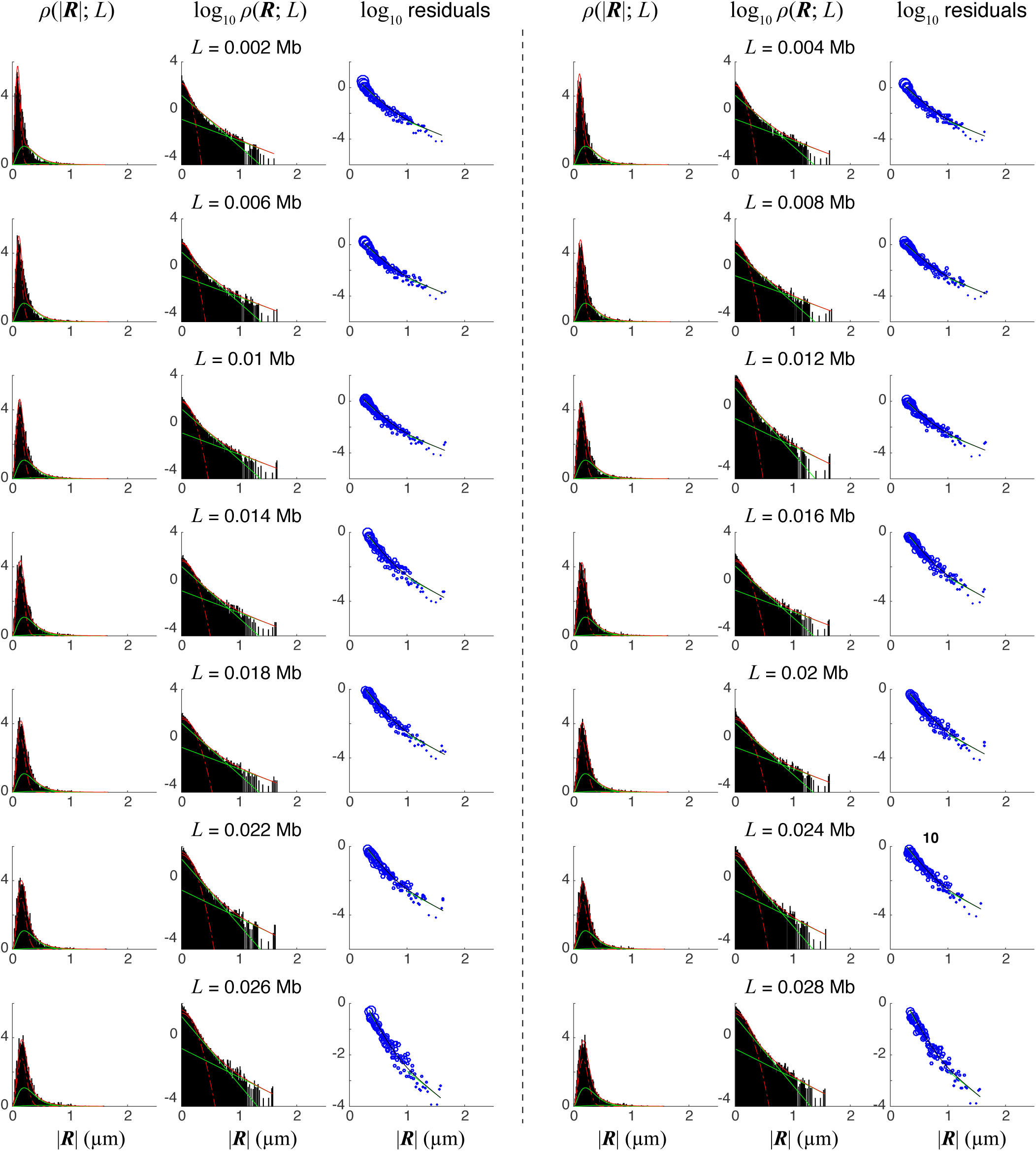
Distance function for ORCA 2 kb reconstructions. Observed probability distribution *ρ* of spatial distances between pairs of loci (blue histograms) overlaid with the model fit (solid red line), at different values of the interlocus distance *L*. See Figure S2 caption for details. We modeled *ρ* as a sum of a Gaussian chain distribution (*p*_*Gauss*_ = 0.540408, |**R**|_*RMS*_ = 0.0025927 *µ*m *L*^0.39361^ for *L* in bp; dotted red line) and two exponential decays in **R** (*p*_*exp*_1 = 0.40666 having decay constant 9.8286 bp^*−*1^, *p*_*exp*1_ = 0.052932 having decay constant 4.1475 bp^*−*1^; green line). The underfit for very small displacements likely comes from the microscope error correction (a constant subtracted term in the squared distance).

**Supplementary Figure S7.**
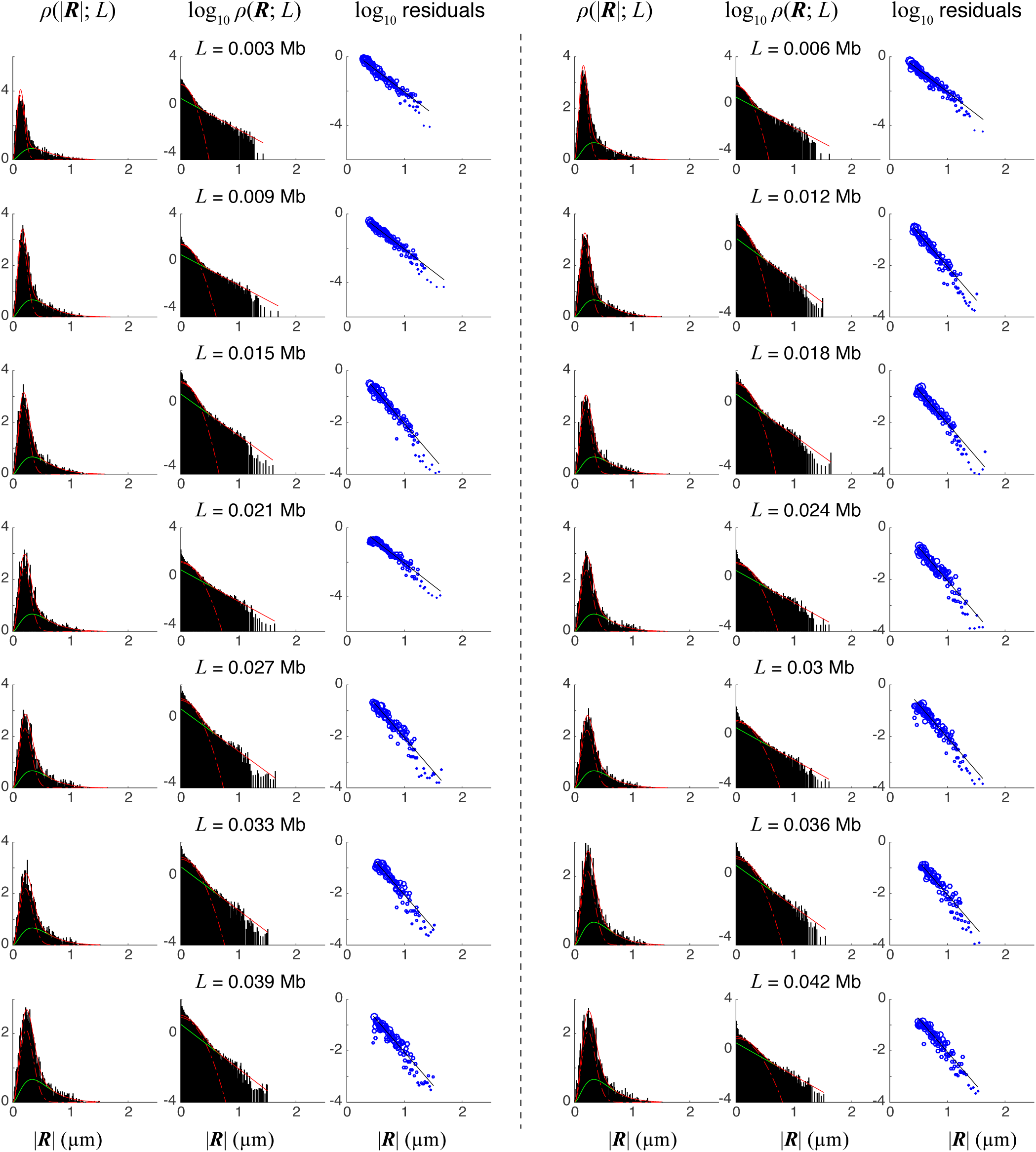
Distance function for ORCA 3 kb reconstructions. Observed probability distribution *ρ* of spatial distances between pairs of loci (blue histograms) overlaid with the model fit (solid red line), at different values of the interlocus distance *L*. See Figure S2 caption for details. We modeled *ρ* as a sum of a Gaussian chain distribution (*p*_*Gauss*_ = 0.58758, |**R**|_*RMS*_ = 0.012231 *µ*m *L*^0.27371^ for *L* in bp; dotted red line) and an exponential decay in |**R**| (*p*_*exp*_ = 0.41242, decay constant 5.9577 bp^*−*^1; green line). This fit differed noticably from that of the 2 kb and 10 kb experiments, showing the importance of calibrating each set of experiments individually.

**Supplementary Figure S8.**
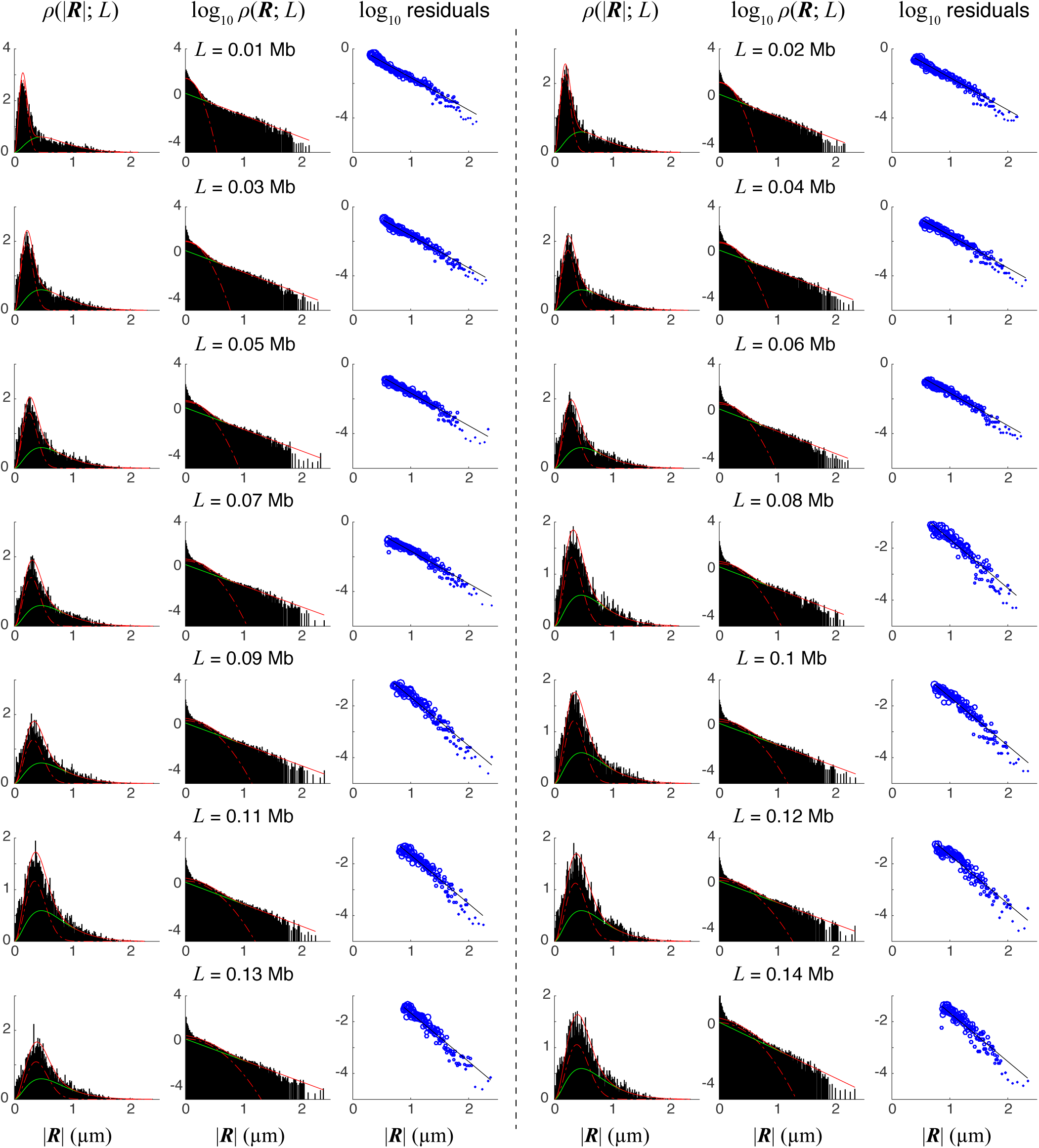
Distance function for ORCA 10 kb reconstructions. Observed probability distribution *ρ* of spatial distances between pairs of loci (blue histograms) overlaid with the model fit (solid red line), at different values of the interlocus distance *L*. See Figure S2 caption for details. We modeled *ρ* as a sum of a Gaussian chain distribution (*p*_*Gauss*_ = 0.49086, |**R**|_*RMS*_ = 0.0025231 *µ*m *· L*^0.42298^ for *L* in bp; dotted red line) and an exponential decay in |**R**| (*p*_*exp*_ = 0.50914, decay constant 4.3206 bp^*−*1^; green line).

**Supplementary Figure S9.**
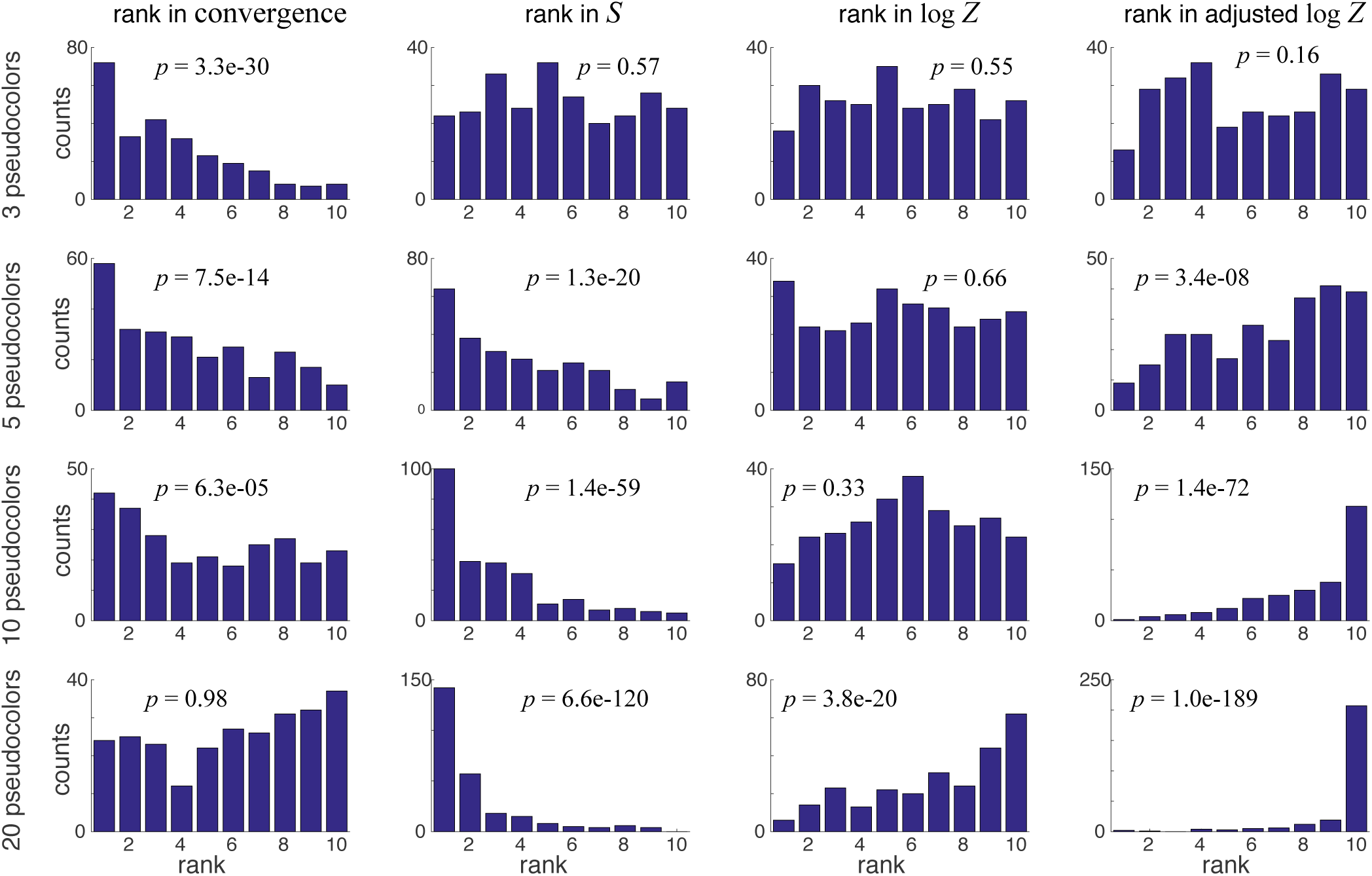
Null hypothesis tests of ORCA 2 kb reconstructions. Ranks of mapping statistics from the true ORCA 2 kb mappings relative to the controls. See Figure S4 caption for details.

**Supplementary Figure S10.**
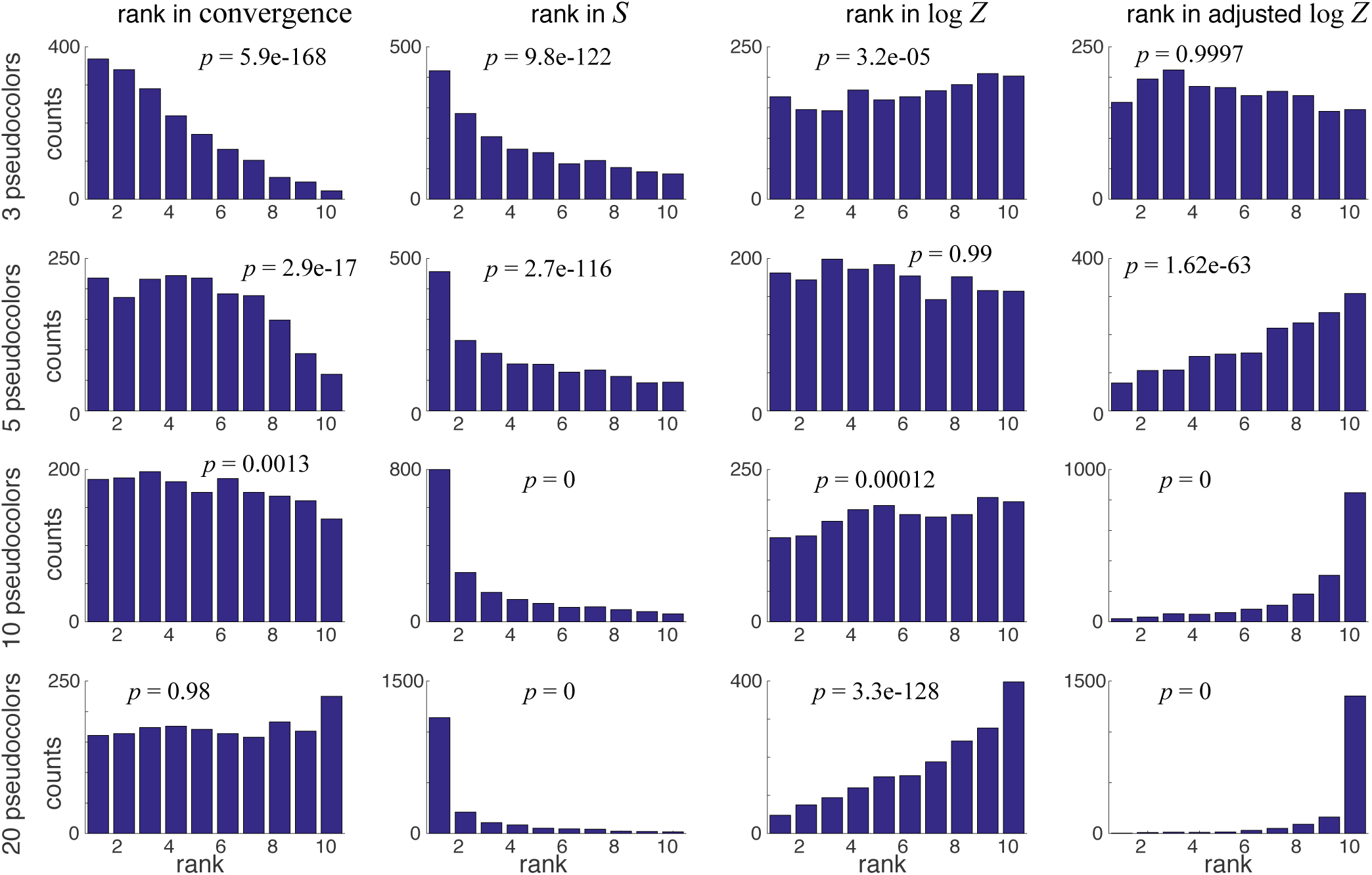
Null hypothesis tests of ORCA 3 kb reconstructions. Ranks of mapping statistics from the true ORCA 3 kb mappings relative to the controls. See Figure S4 caption for details.

**Supplementary Figure S11.**
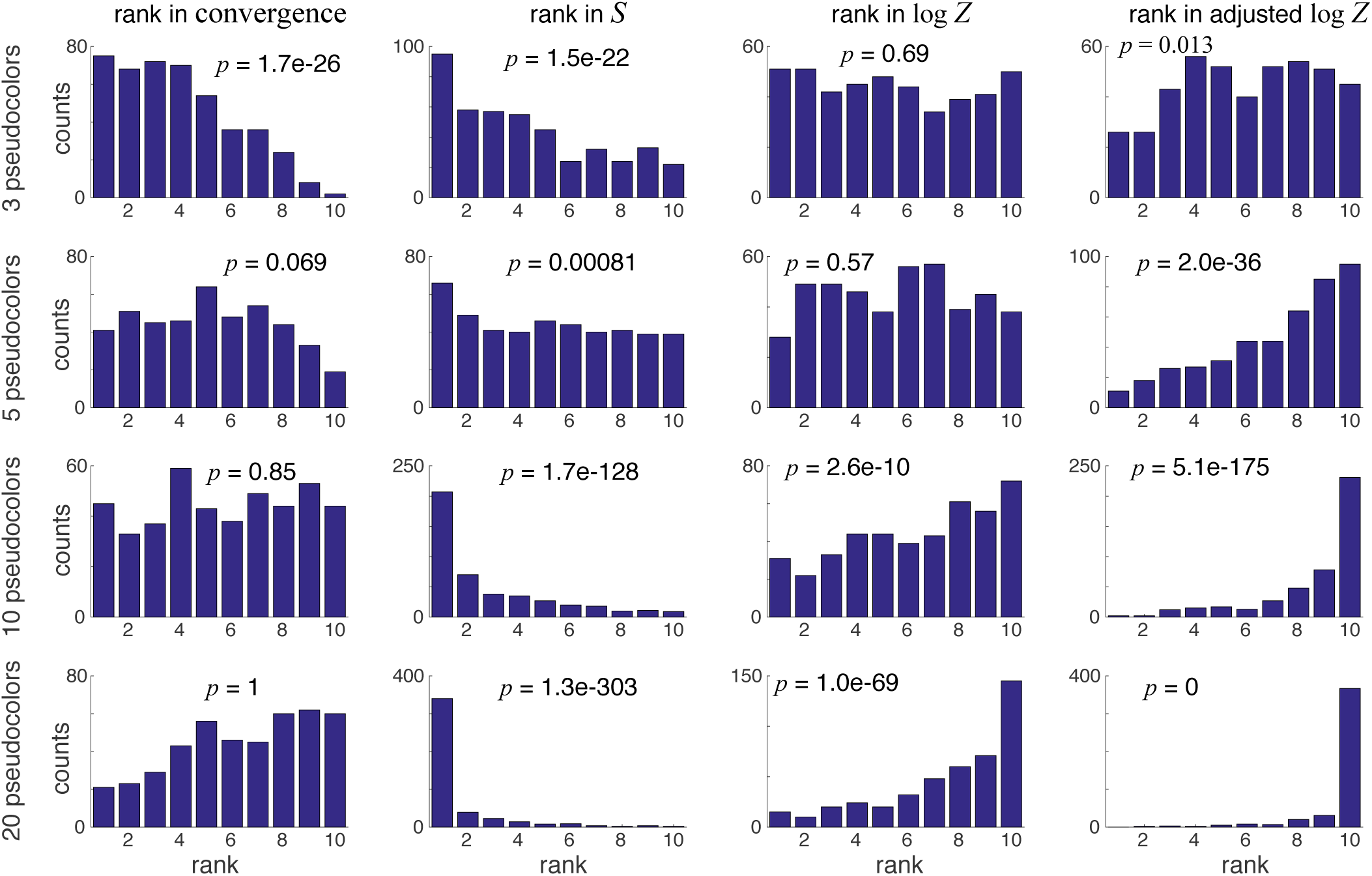
Null hypothesis tests of ORCA 10 kb reconstructions. Ranks of mapping statistics from the true ORCA 10 kb mappings relative to the controls. See Figure S4 caption for details.

**Supplementary Figure S12.**
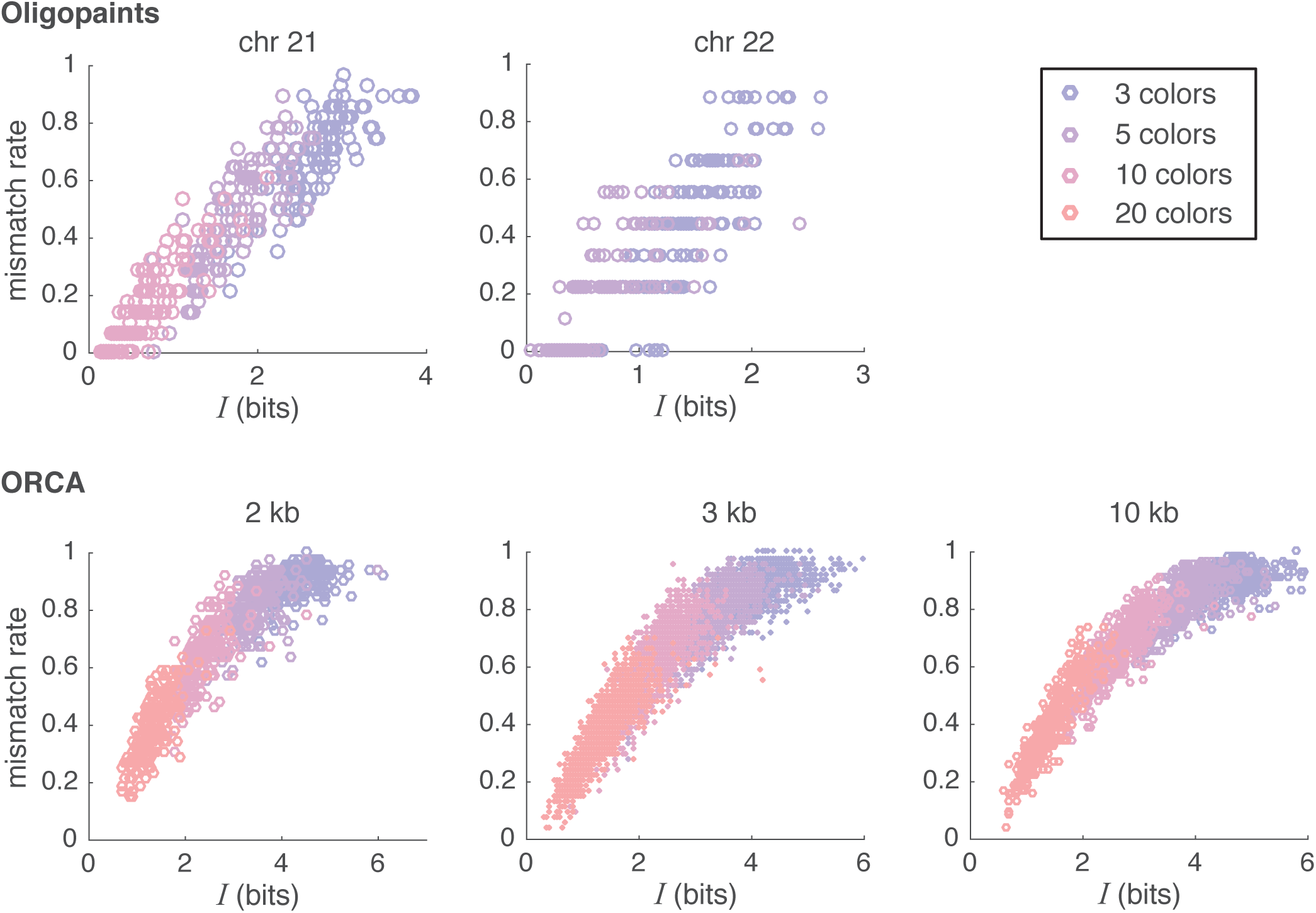
Spot-assignment error rate vs. unrecovered information per locus *I*. This plot connects the abstract measure of information recovery *I*, which is defined over mapping probabilities generated by `align3d`, to the more tangible error rate when trying to identify imaged spots, by inferring an explicit conformation from the strongest mapping probabilities. The various experiments produce similar curves, indicating a simple universal relation between the two measures.

**Supplementary Figure S13.**
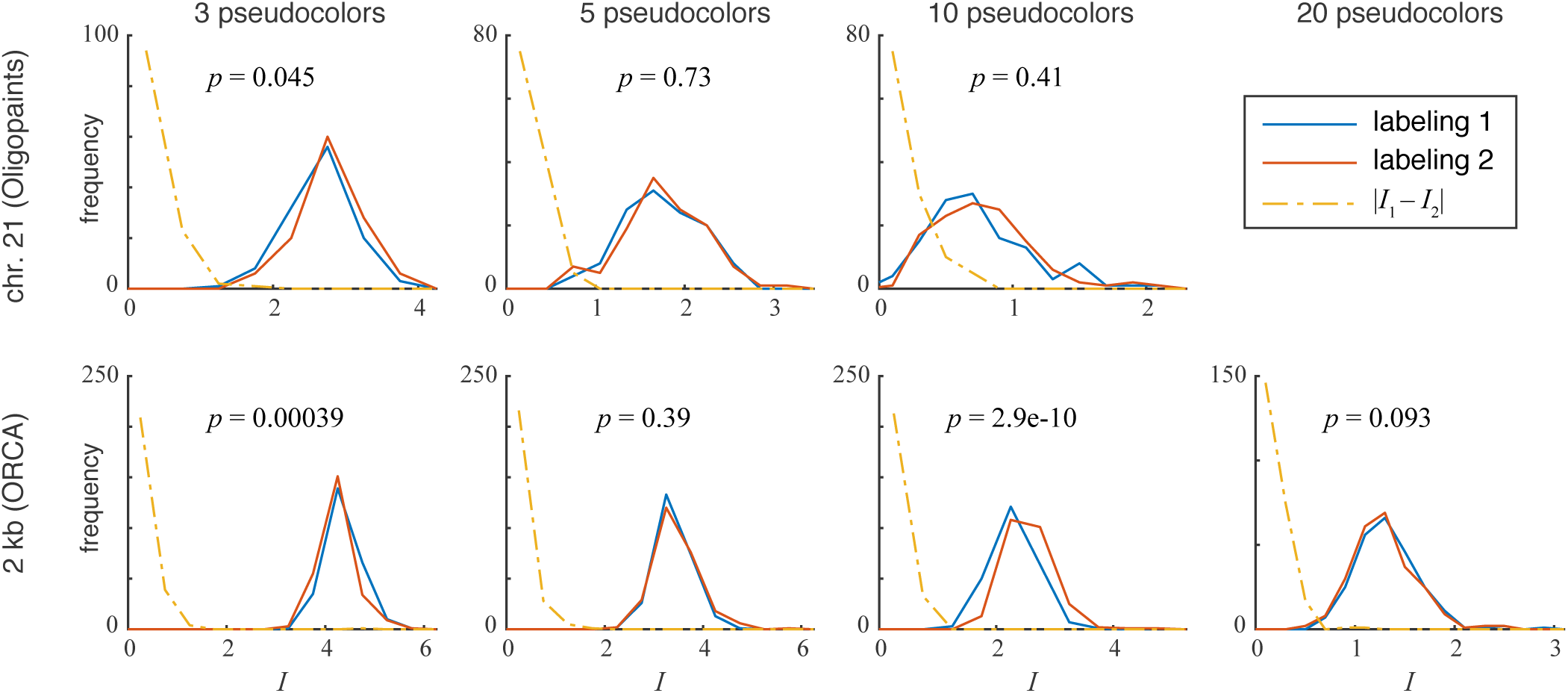
Reconstruction quality from independent label coloring. Each plot compares histograms of unrecovered information *I* from two independent, randomly-generated color assignments for the labeled loci and the corresponding imaged spots. Only one experiment showed a sizeable difference between the two labeling patterns, and the difference was small although statistically robust (see *p*-values on plot). For comparison, the dot-dash line is a histogram of the chromosome-by-chromosome difference in *I* for the respective experiment. These experiments seem to indicate that the quality of a randomly-chosen coloring pattern mainly varies randomly between chromosomes, and that the systematic quality difference between labeling patterns is fairly small. However, we only tested a few labeling patterns.

**Supplementary Figure S14.**
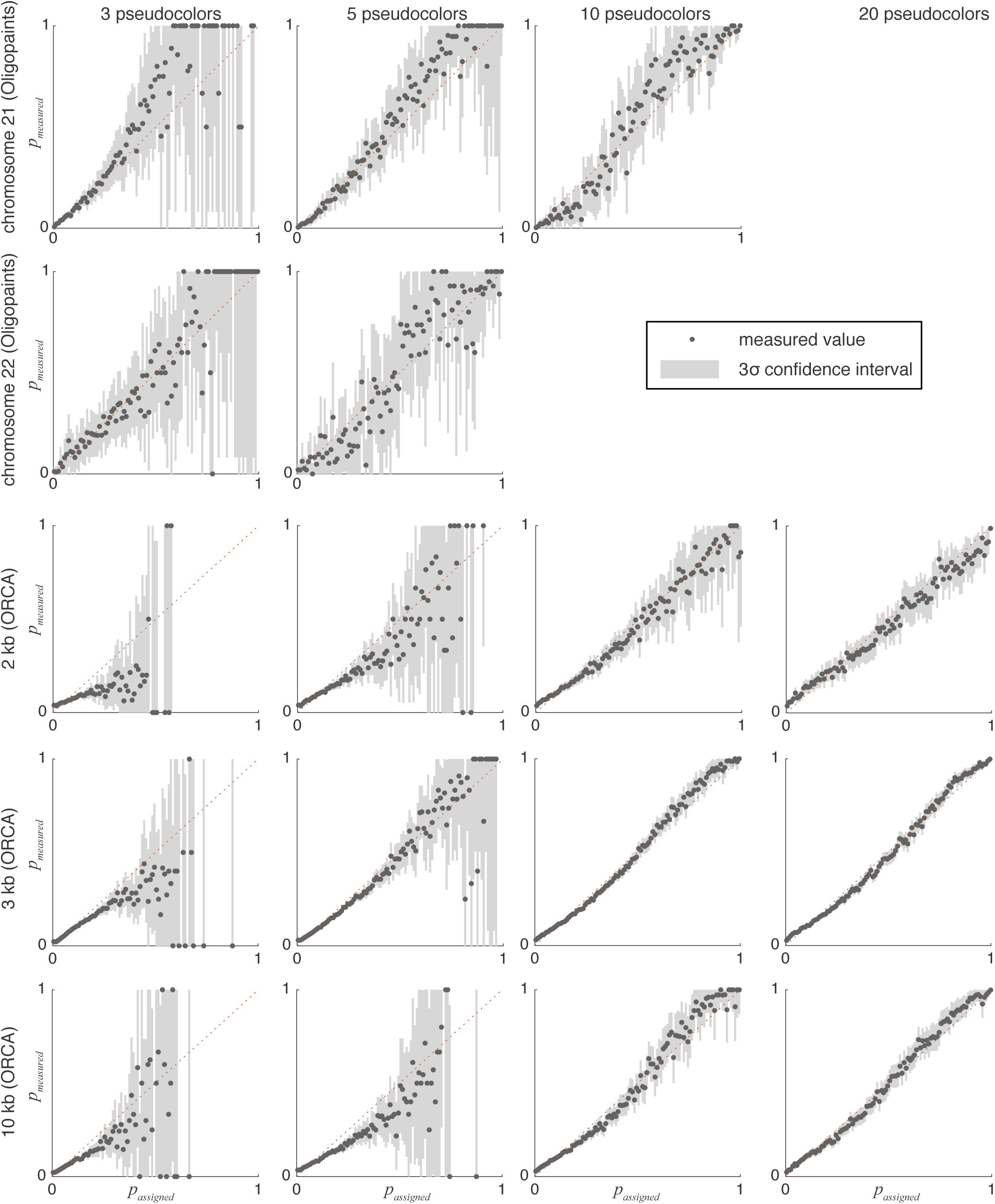
Assigned mapping *p*-values versus their likelihoods of being true mappings. Mapping *p*-values are split into 100 bins (x axis), and the fraction of true mappings in each bin are plotted using a dot on the y axis. Grey shaded regions show the 3*σ* confidence interval owing to counting statistics. Ideal mapping *p*-values would lie along the *p*_*measured*_ = *p*_*assigned*_ diagonal (dotted line).

## Footnotes

1 Ref. [18] argues that reconstruction quality depends on a competition between the number of pseudocolors and the *density* (not number) of spots in the image, and that the number of pseudocolors required for near-perfect results is on the order of the number of competing loci that are spatially as close to a given locus *L*_*i*_ as either of its neighbors *L*_*i−*1_ or *L*_*i*+1_.

